# Evolutionary rate covariation analysis of E-cadherin identifies Raskol as regulator of cell adhesion and actin dynamics in *Drosophila*

**DOI:** 10.1101/429472

**Authors:** Qanber Raza, Jae Young Choi, Yang Li, Roisin M. O’Dowd, Simon C. Watkins, Yang Hong, Nathan L. Clark, Adam V. Kwiatkowski

## Abstract

The adherens junction couples the actin cytoskeletons of neighboring cells to provide the foundation for multicellular organization. The core of the adherens junction is the cadherin-catenin complex that arose early in the evolution of multicellularity to link cortical actin to intercellular adhesions. Over time, evolutionary pressures have shaped the signaling and mechanical functions of the adherens junction to meet specific developmental and physiological demands. Evolutionary rate covariation (ERC) identifies genes with correlated fluctuations in evolutionary rate that can reflect shared selective pressures and functions. Here we use ERC to identify genes with evolutionary histories similar to *shotgun (shg)*, which encodes the Drosophila E-cadherin (DE-Cad) ortholog. Core adherens junction components α-catenin and p120-catenin displayed strong ERC correlations with *shg, indicating that they evolved under similar selective* pressures during evolution between Drosophila species. Further analysis of the *shg* ERC profile revealed a collection of genes not previously associated with *shg* function or cadherin-mediated adhesion. We then analyzed the function of a subset of ERC-identified candidate genes by RNAi during border cell (BC) migration and identified novel genes that function to regulate DE-Cad. Among these, we found that the gene *CG42684*, which encodes a putative GTPase activating protein (GAP), regulates BC migration and adhesion. We named *CG42684 raskol* (“to split” in Russian) and show that it regulates DE-Cad levels and actin protrusions in BCs. We propose that Raskol functions with DE-Cad to restrict Ras/Rho signaling and help guide BC migration. Our results demonstrate that a coordinated selective pressure has shaped the adherens junction and this can be leveraged to identify novel components of the complexes and signaling pathways that regulate cadherin-mediated adhesion.

**Author Summary:** The establishment of intercellular adhesions facilitated the genesis of multicellular organisms. The adherens junction, which links the actin cytoskeletons of neighboring cells, arose early in the evolution of multicellularity and selective pressures have shaped its function and molecular composition over time. In this study, we used evolutionary rate covariation (ERC) analysis to examine the evolutionary history of the adherens junction and to identify genes that coevolved with the adherens junction gene *shotgun, which encodes the Drosophila E-cadherin (DE-Cad). ERC analysis of shotgun* revealed a collection of genes with similar evolutionary histories. We then tested the role of these genes in border cell migration in the fly egg chamber, a process that requires the coordinated regulation of cell-cell adhesion and cell motility. Among these, we found that a previously uncharacterized gene *CG42684*, which encodes a putative GTPase activating protein (GAP), regulates the collective cell migration of border cells, stabilizes cell-cell adhesions and regulates the actin dynamics. Our results demonstrate that components of the adherens junction share an evolutionary history and that ERC analysis is a powerful method to identify novel components of cell adhesion complexes in *Drosophila*.

## Introduction

The adherens junction (AJ) is a multiprotein complex that is essential for intercellular adhesion in metazoa. The core of the AJ is the cadherin-catenin complex. Classical cadherins are single-pass transmembrane proteins with an extracellular domain that mediates calcium-dependent homotypic interactions. The adhesive properties of classical cadherins are driven by the recruitment of cytosolic catenin proteins to the cadherin tail: p120-catenin binds to the juxta-membrane domain and β-catenin binds to the distal part of the tail. β-Catenin recruits α-catenin to the cadherin-catenin complex. α-Catenin is an actin-binding protein and the primary link between the AJ and the actin cytoskeleton [1–3].

The primary function of the AJ is to link actin to intercellular junctions. It is believed the AJ arose early in the evolution of multicellular metazoans to coordinate epithelial tissue formation and organization [4–7]. The AJ has since evolved to function in a range of physiological and developmental processes, including cell polarity, collective cell migration and cell division [8, 9]. AJ function in these diverse processes requires an array of ancillary regulatory proteins, including kinases, signaling molecules and adaptor proteins [10–14]. Defining the molecular networks that regulate AJ biology is critical to understanding cadherin-mediated adhesion in normal and disease states.

Evolutionary rate covariation (ERC) analysis is a comparative genomic approach that has been used successfully to identify genes with shared functions in canonical protein complexes and biological processes in prokaryotes, fungi, Drosophila and mammals [15–21]. ERC analysis takes advantage of the evolutionary rates shared between co-functional genes that have correlated rates of change due to common selective pressures. ERC represents the correlation coefficient of branch-specific evolutionary rates that are estimated from the phylogenetic trees of a pair of genes and their orthologs from multiple species[19]. ERC analysis permits the identification of genes within a given genome that evolved in a correlated manner and hence might function in the same pathway or molecular complex. These genes can then be screened by RNAi-based knockdown or similar genetic approaches to validate their role in a relevant biological process.

Border cell (BC) migration in the developing *Drosophila* egg chamber requires coordinated cell adhesion and migration. During BC migration, a group of 6–8 follicular cells delaminate from the anterior most tip of the epithelium and undergo haptotaxis and migrate collectively towards the developing oocyte [22, 23]. The BC cluster consists of migratory BCs and a centrally positioned pair of polar cells (PCs) that signal to BCs and contribute to cluster adherence [23]. BC migration is highly dependent on *Drosophila E-*Cadherin (DE-Cad, encoded by *shotgun (shg*)) [24–26]. Upregulation of DE-Cad is essential for the initial delamination and subsequent migration of BC since disruption of DE-Cad-mediated adhesion affects the ability of BC to detach from the follicular epithelium (FE) and collectively migrate [24, 25].

We performed ERC analysis of *shg* to identify genes that share a common evolutionary history, and therefore may share an overlapping function, with shg and assessed their role in BC migration. Genes encoding the primary components of AJ, including *α-Cat* and *p120-catenin*, display high ERC values relative to shg and one another, suggesting that these genes and their protein products are co-functional, which is well described in the literature [1, 27]. We show that genes showing high ERC values with *shg* are enriched for membrane-associated proteins and proteins that function in E-cadherin-dependent biological processes. We then conducted an RNAi-mediated genetic screen in BCs with 34 high-ranking ERC candidates and identified both novel and known genes that function to regulate DE-Cad at cell contacts. Among those, we characterized a GTPase activating protein (GAP) domain encoding gene, *CG42684*, which we have named “*raskol*” after the Russian term “to split”. We show that Raskol colocalizes with DE-Cad, regulates DE-Cad levels at the BC-BC interface and modulates actin-rich protrusions during BC migration. Our results demonstrate that components of the AJ share an evolutionary history and that ERC analysis is a powerful method to identify novel components of cell adhesion complexes in *Drosophila*.

## Results

### ERC analysis identifies genes that coevolved with *shg*

We used ERC analysis to identify novel genes that regulate DE-Cad-mediated adhesion in *Drosophila*. Since ERC signatures are often observed between proteins that function in a molecular complex [15, 16, 18, 19], we evaluated the ERC values of the fly AJ components – shotgun (*shg*, DE-Cad), *armadillo* (*arm*; β-catenin in vertebrates), *α-catenin (α-Cat*), *p120-catenin* (*p120ctn), Vinculin* (*Vinc*) and canoe (*cno*, afadin in vertebrates). Notably, *shg, α-Cat* and *p120ctn* displayed positive ERC values with each other (Fig 1A). *arm, Vinc* and *cno* did not show elevated ERC values relative to *shg, α-Cat* or *p120ctn*. Since Arm, Vinc and Cno are known to function independently of cadherin-mediated adhesion [28–31], we speculate that alternative selective pressures have influenced the evolution of these genes in flies, likely obscuring any ERC signature with purely AJ genes. Nonetheless, ERC analysis suggested that the AJ components DE-Cad, α-Cat and p120ctn coevolved to maintain their collective function in cell-cell adhesion. We postulated that other genes whose products regulate AJ biology would have similar evolutionary histories to maintain functionality.

| Genes | ERC value | Function in flies | Ortholog(s) (score*) | Ortholog function in mammals |
| --- | --- | --- | --- | --- |
| *grk* | 0.62 | Growth factor (EGF) | Nrg (1/15) | EGF family receptor ligand |
| *cac* | 0.62 | Voltage-gated Ca<sup>2+</sup> channel subunit | Cacna1 (11/15) | Voltage-gated Ca<sup>2+</sup> channel subunit |

We then used ERC analysis to identify genes with high ERC values relative to the core of the AJ complex, *shg* (S1 Table). We identified 137 genes with ERC values of 0.4 or greater, representing the top 1.3% of *shg* ERC values. Since α-*Cat* has an ERC value of 0.47 relative to *shg* (Fig 1A), placing their ERC value in the top 0.6 % of all gene pairs, we reasoned that genes with similar or higher ERC values would represent genes with similar evolutionary histories to shg. Accordingly, the thresholded *shg* ERC list contains genes with described roles in AJ regulation such as *Hrb98DE* [32, 33], *PDZ-GEF* [34, 35], *babo* [36, 37], *CG16952 [*38, 39] and *Rab5* [40–42] (Table 1). Excitingly, the majority of identified genes have not been associated with the AJ and include transcription factors, kinases, GTPase regulatory proteins and calcium channel regulators (Table 1). A previous genomic RNAi screen conducted in *Drosophila* S2 cells [43] and E-Cadherin proximity biotinylation screens in epithelial cells [44, 45] identified multiple hubs of interactors and regulators. Cross referencing the *shg* ERC list with the hits from these screens revealed only a few common genes such as *RhoGAPp190, Rab5, Appl* and *Stim*. This suggests that ERC analysis is identifying additional shg regulatory components that were undetected in genetic or proteomic screens.

**Table 1.**
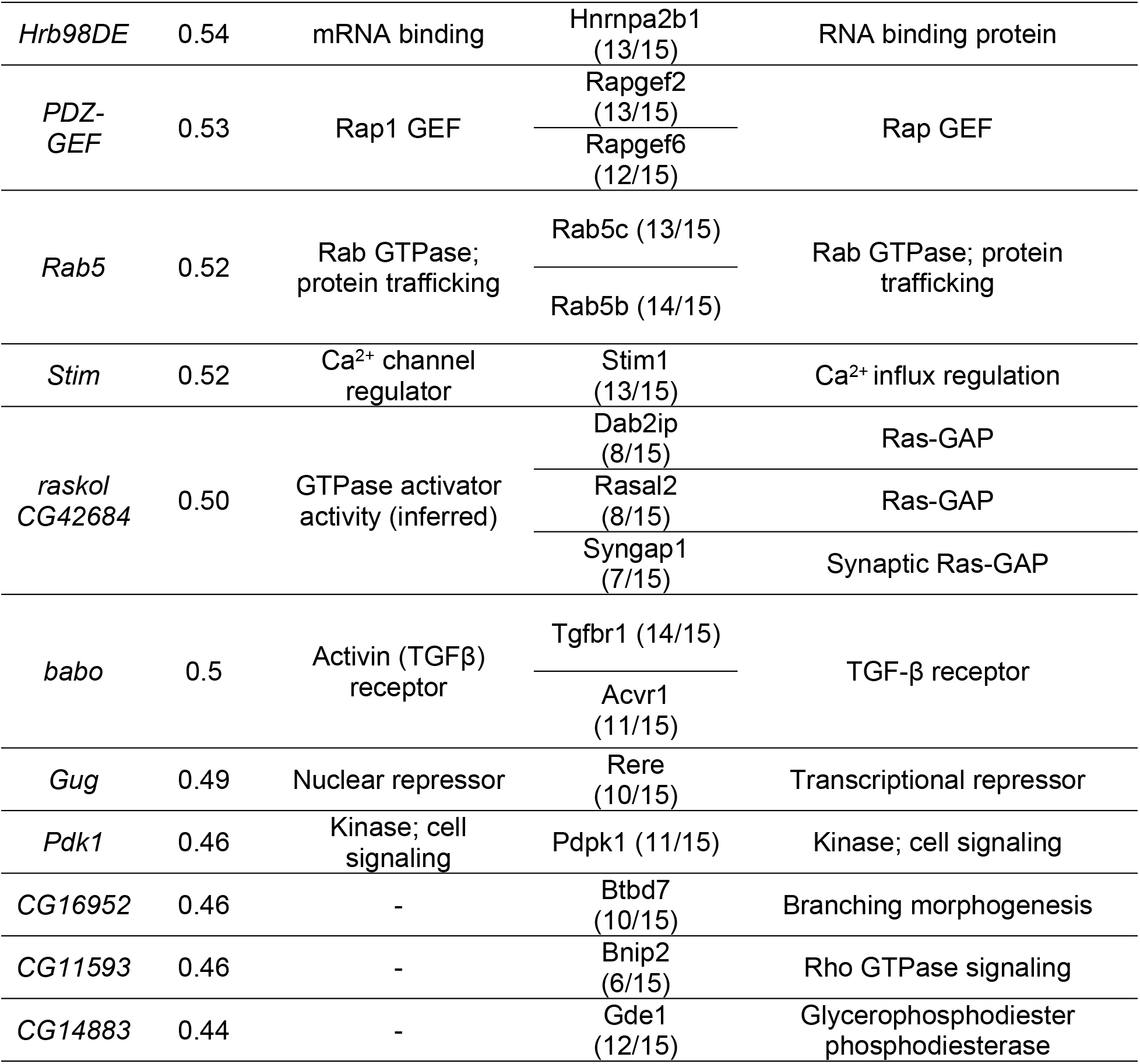
Function of selected conserved genes identified in shg ERC analysis. *Score from flybase (www.flybase.org) orthologue database. Ratio indicates sequence alignment algorithms that reported significant homology with mammalian orthologues.

Using gene ontology (GO) based enrichment analysis, we found that the *shg* ERC genes are enriched for plasma membrane (PM) localized proteins and PM-associated protein complexes (Fig 1B). Additionally, genes that function in biological processes requiring E-cadherin-mediated adhesion, such as wing disc morphogenesis, imaginal disc morphogenesis, epithelial morphogenesis, cell migration and AJ organization, were significantly overrepresented in the *shg ERC list (Fig 1C). Next, we* analyzed the molecular functions of the human orthologs of the ERC identified genes. We found that genes involved in epithelial AJ remodeling and cancer molecular mechanism were overrepresented (Fig 1D; S2 and S3 Table). Also, genes that function in RhoA, CCR5, TGF-β, PTEN and AJ-mediated signalling pathways were enriched (Fig 1D, S2 and S3 Table). Thus, the *shg* ERC list contains genes with established roles in regulating AJs as well as novel candidate genes.

**Fig 1.**
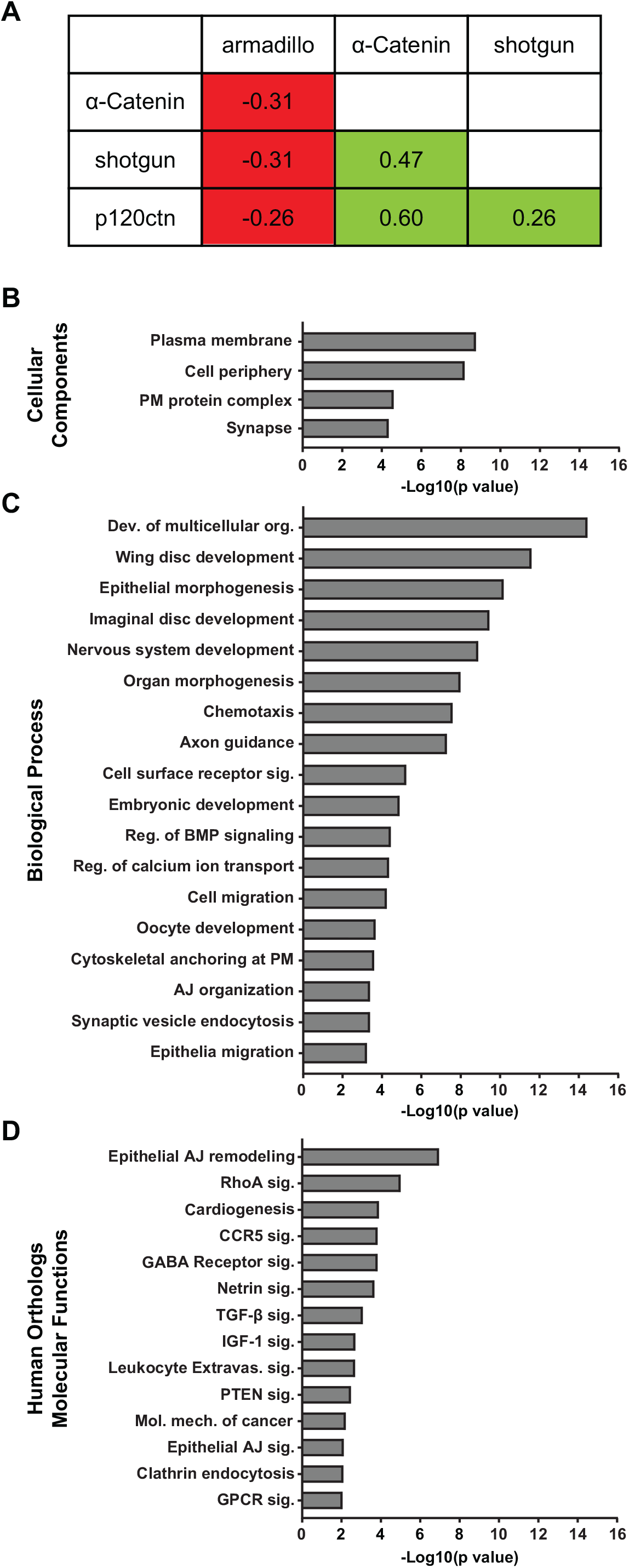
Fig 1. Gene ontology enrichment analysis for genes identified in *shg* ERC analysis. A. ERC analysis reveals an evolutionary relationship between *shg, α-cat* and *p120ctn*. B. Genes from the *shg* list with ERC values ≥ 0.4 are enriched for genes that encode plasma membrane and cell periphery proteins. C. *shg* ERC list is enriched for genes with established roles in regulating biological processes that require cadherin-mediated adhesion. D. Human orthologs of the *shg* ERC list are involved in remodeling of epithelial AJs and multiple canonical pathways including RhoA, TGF-β, PTEN and AJ signaling.

### RNAi screen in BCs identifies genes that regulate cell-cell adhesion

To evaluate the function of genes in the *shg* ERC list, we conducted an *in vivo* RNAi-based genetic screen in the *Drosophila* egg chamber. We analyzed BC collective cell migration (CCM) because it is regulated by DE-Cad and is a powerful system to study the interplay between cell migration and cell-cell adhesion (Fig 2A) [23, 24]. BC detachment from the FE and concomitant CCM requires increased DE-Cad expression and loss of DE-Cad arrests BC migration [24, 25].

**Fig 2.**
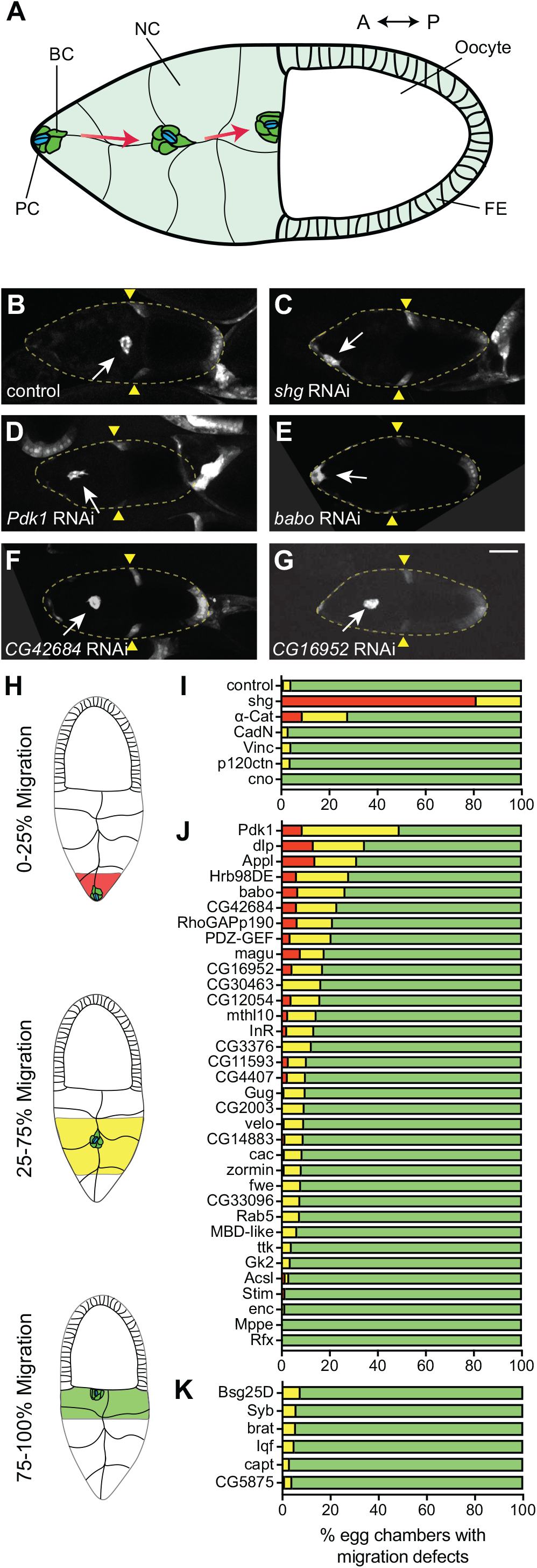
*shg* ERC genes regulate BC migration. A. Cartoon representation of BC migration during egg chamber development. BC, border cell; PC, polar cell; NC, nurse cell; FE, follicular epithelium; A, anterior; P, posterior. B-G. Representative images of egg chambers expressing UAS-GFP and UAS-RNAi for control (luciferase) (B), shg (C), *Pdk1* (D), *babo (E), CG42684* (F) or CG16952 (G) under the control of *slbo-GAL4*. White arrow indicates BC cluster position. Yellow triangles mark FE retraction border. Maximum projections of 20 μm z-stacks are shown. Posterior is to the right in all images. H. Border cell migration scoring classes. I-K. Percentage of egg chambers in each class displaying migration defects in AJ-related genes (I), *shg* ERC target genes (J) and random negative control genes (K). Red 0–25%, yellow 25–75% and green 75–100%. N values are listed in S5 Table. Scale bar in G is 50 μm and applies to B-G.

First, we downregulated levels of individual AJ genes by using the GAL4/UAS system to drive UAS-RNAi transgenes in the migrating BC cluster. We used a BC specific driver, *slowborder-*GAL4 (*slbo*-GAL4) [46, 47] to drive expression of a UAS-GFP reporter and a UAS-RNAi transgene targeted against the gene of interest. Stage 10 egg chambers expressing RNAi and GFP in BCs were fixed and scored for BC cluster position along the anterior-posterior migration axis (Fig 2H). In control egg chambers, nearly all BC clusters completed migration and were positioned adjacent to the oocyte (Fig 2B and 2I). In contrast, downregulation of *shg* caused a BC migration failure or delay in all egg chambers (Fig 2C and 2I). BC migratory defects were less severe in egg chambers with reduced expression of *α-Cat* compared to shg; however, the prevalence of defects was higher than in control egg chambers (Fig 2I). We could not assess the effect of *arm* downregulation since *arm* RNAi expressing flies did not survive to adulthood. The downregulation of CadN, Vinc, p120ctn or *cno* did not lead to BC migratory defects (Fig 2I).

Next, we screened 34 genes from the *shg* ERC list. We focused on genes that are expressed in the ovary [48] and for which an RNAi stock was readily available (S4 Table). The downregulation of target genes displayed a variable range of BC migration defects with 12 genes displaying defects in more than 15% of egg chambers assessed compared to 4% in control (Fig 2J). We also randomly selected and screened six genes that had either very low ERC values or did not appear in the *shg* ERC list for migration defects. As expected, we did not observe strong migration defects when these genes were knocked down (Fig 2K). Moreover, the shg ERC genes showed statistically lower average migration than the random negative control genes (Wilcoxon rank sum test, p=0.0162), supporting the hypothesis that genes with correlated evolutionary histories share functional characteristics. *Pdk1* knockdown resulted in the most penetrant phenotype with 50% of egg chambers displaying either a failure or delay in BC migration (Fig 2D and 2J). Knockdown of *babo* and *CG42684* caused migration delays similar to *α-Cat* (Fig 2E–J). Additionally, knockdown of multiple genes resulted in considerable migration delays relative to the control including *CG16952* (Fig 2D), *Hrb98DE*, magu, *InR, RhoGAPp190, PDZ-GEF* and *CG11593 (Fig 2J). Conversely*, knockdown of genes such as *enc, mppe and rfx* did not affect BC migration (Fig 2J).

While screening *α-Cat* knockdown egg chambers, we noticed that about 20% of the egg chambers displayed a cluster disassociation phenotype where one or more BCs had separated from the cluster (Fig 3B and 3G). Since this phenotype is indicative of cell adhesion defects in BCs [24], we scored cluster disassociation for all genotypes. In control BCs expressing luciferase RNAi, disassociated clusters were rarely observed (Fig 3A and 3G). Likewise, downregulation of genes with low ERC values did not show a cluster disassociation phenotype (Fig 3I). Knockdown of *CadN, vinc, p120ctn* and *cno* also did not cause a penetrant cluster disassociation phenotype (Fig 3G). However, cluster disassociation phenotypes were observed with a number of ERC target genes. *CG42684, PDZ-GEF, CG16952, CG11593, zormin, cac* and *Rab5* displayed similar or higher cluster disassociation phenotypes compared to *α-Cat* (Fig 3). Overall, these ERC target genes, chosen for their high *shg* ERC values, exhibited the cluster disassociation phenotype significantly more often than genes with low *shg* ERC values (Wilcoxon rank sum test, p= 0.00022). Together, these results demonstrate that highly ranked genes in the *shg* ERC list contain factors that may regulate cell adhesion during BC collective cell migration.

**Fig 3.**
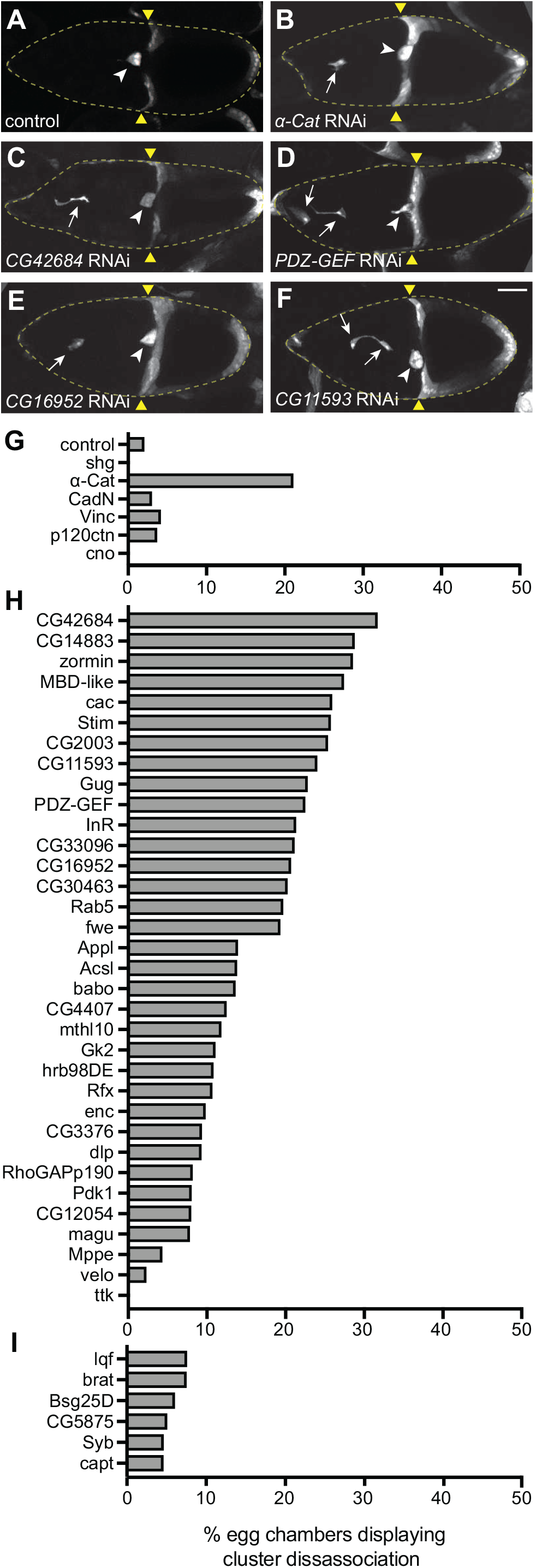
*shg* ERC genes maintain BC adhesion during migration. A-F. Egg chambers expressing UAS-GFP and UAS-RNAi for control (A), *α-Cat* (B), *CG42684* (C), PDZ-GEF (D), *CG16952 (E) or CG11593* (F). White arrows mark disassociated BCs. White arrowheads mark BC cluster adjacent to oocyte. Yellow triangles mark the FE retraction border. Maximum projections of 20 μm z-stacks are shown. G-I. Percentage of egg chambers displaying a cluster disassociation phenotype in AJ genes (G), *shg* ERC target genes (H) and random negative control genes (I). Scale bar in F is 50 μm and applies to A-F.

### Top ERC candidates regulate DE-Cad levels in BCs

Since knockdown of a subset of *shg* ERC list genes disrupted BC migration, we hypothesized that these genes might regulate DE-Cad at BC contacts. To test this, we quantified DE-Cad levels along cell-cell contacts between BCs in RNAi-expressing clusters. We used a DE-Cad-GFP knock-in stock that express DE-Cad at endogenous levels [49] and drove RNAi constructs under the control of *slbo*-GAL4. In control egg chambers where either luciferase RNAi or RFP were expressed, high DE-Cad levels were observed at BC-BC contacts (Fig 4A, S2A Fig). As expected, *shg* RNAi expression severely reduced DE-Cad levels in the BC cluster (Fig 4B). Knockdown of *CG42684* (Fig 4C), *CG16952* (Fig 4D), *CG11593* (Fig 4E), *babo (Fig 4G) and Hrb98DE* (S2B Fig) caused a significant reduction in DE-Cad levels at BC-BC contacts. Interestingly, knockdown of *Pdk1*, a kinase in the PI3K pathway [50], did not affect DE-Cad levels in BCs even though it displayed the most prominent migration defect (Fig 4F). As expected, expression of RNAi for genes capt and *CG5872*, which have very low ERC values relative to *shg* and were therefore not predicted to operate in this pathway, did not cause a reduction in DE-Cad levels at BC-BC contacts (S2C and S2D Fig). Collectively, these results indicate that a significant subset of genes with high ERC values relative to *shg*, many of which were previously not associated with cadherin function, regulate DE-Cad levels at BC-BC contacts.

**Fig 4.**
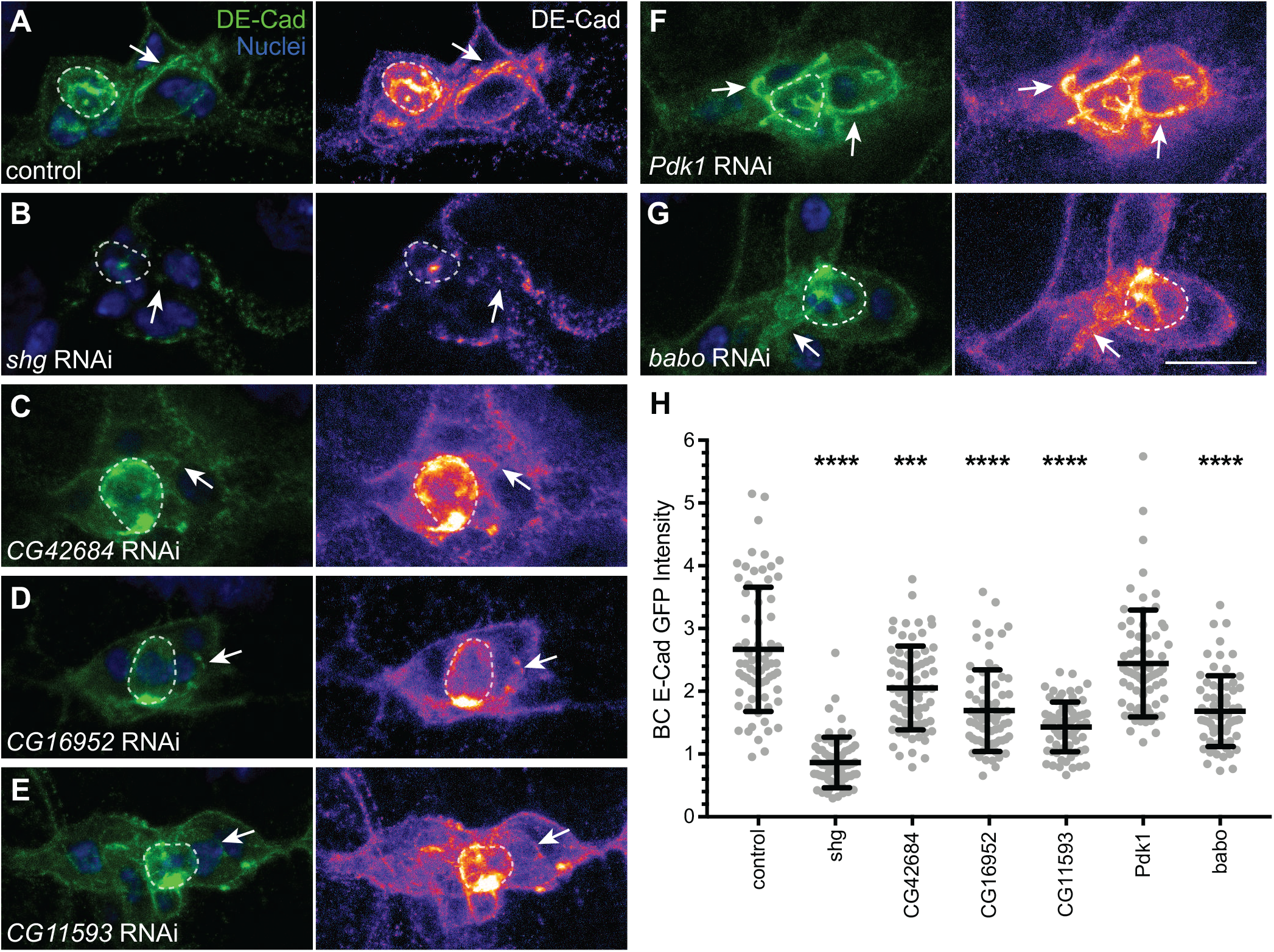
*shg* ERC genes regulate DE-Cadherin levels in BCs. A-G. Representative images of BC clusters stained for nuclei (blue) and expressing DE-Cad-GFP and control (A,), shg (B), CG42684 (C), *CG16952 (D), CG11593* (E), Pdk1 (F) or *babo* (G) RNAi under the control of *slbo*-GAL4. White arrows mark BC-BC contacts. PCs are outlined by a dashed line. Maximum projections of 5 μm z-stacks are shown. H. Quantification of DE-Cad levels at BC-BC contacts in control (n=69), *shg* (n=66), *CG42684 (n=72), CG16952* (n=69), *CG11593*(n=69), *Pdk1*(n=66) and babo (n=69) RNAi-expressing BCs. **** – p<0.0001, *** – p<0.001. Error bars represent mean ± SD. Scale bar in G is 10 μm and applies to A-G.

### Raskol colocalizes with DE-Cadherin

Knockdown of *CG42684* displayed the most severe cell disassociation phenotype amongst all genes tested (Fig 3). CG42684 is reported to localize at the cell cortex and is enriched specifically at the apical surface of epithelial cells in *Drosophila* embryo [51] though almost nothing is known about its molecular function in flies. The mammalian orthologs of CG42684, Rasal2 and Dab2IP, also localize to the PM [52]. Interestingly, and similar to the impact we report here for CG42684, downregulation of Rasal2 disrupts E-Cadherin localization at the cell contacts [53], though the mechanism for this disruption remains poorly defined and its conservation across phyla has yet to be reported. Therefore, we wanted to determine whether CG42684, which we named “Raskol” (Russian for “to split”) associates with DE-Cad along the cell membrane. Consistent with earlier localization studies, a YFP-trap stock expressing Raskol-YFP localized to the cell periphery in embryonic epidermal cells (S3 Fig) [51]. We used this stock to assess Raskol colocalization with DE-Cad along the BC membrane. In *Drosophila* stage 8 embryos, before BCs have delaminated, Raskol localized to the cell membrane of BCs and PCs and was enriched at the PC apical membrane (Fig 5A, S2B Fig). During migratory stages, Raskol localization persisted at the PC apical membrane and at the cell-cell contacts of BCs and PCs (Fig 5B). Immunolabeling of egg chambers with DE-Cad antibody or imaging of endogenously RFP-tagged DE-Cad revealed strong colocalization between DE-Cad and Raskol (R=0.63, n=15) at the apical surface of PCs and at BC-BC contacts (Fig 5C and 5D). We also observed colocalization in the FE, particularly along the apical membrane (Fig 5A and 5C; S2A and S2B Fig). Raskol was not present at cell contacts between NCs (S2B Fig).

**Fig 5.**
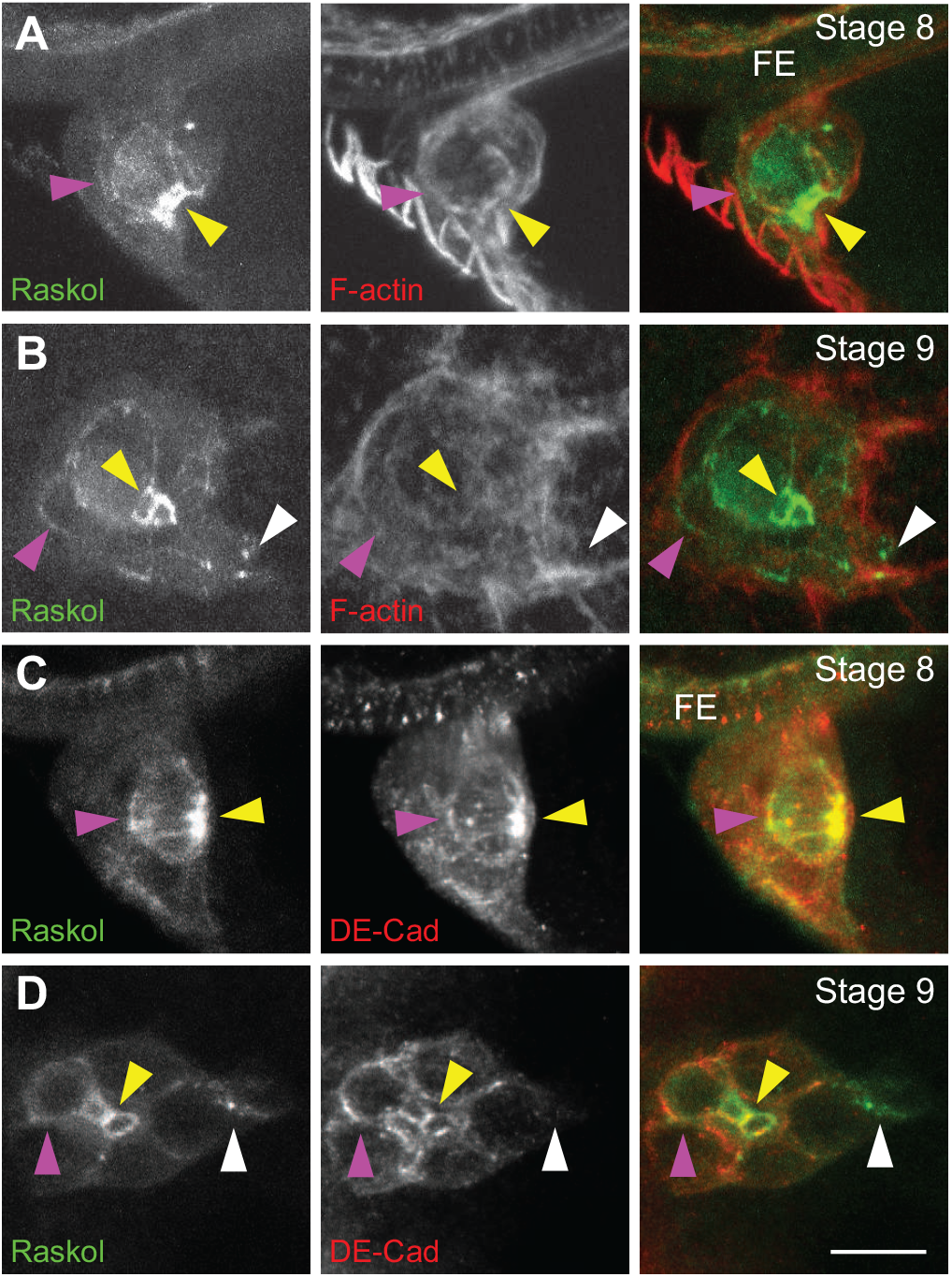
Raskol localizes to BC contacts. A-B. Egg chambers expressing Raskol-YFP and stained for F-actin (red) at stage 8 (A) and stage 9 (B). C-D. Egg chamber expressing Raskol-YFP and stained with DE-Cad (red) at stage 8 (C) and stage 9 (D). Yellow triangles indicate the apical side of PCs. Purple triangles mark cell-cell contacts. White triangles mark Raskol localization in the leading protrusion. FE – follicular epithelium. Maximum projections of 5 μm z-stacks are shown. Scale bar in D is 10 μm and applies to A-D.

Next, we determined whether Raskol colocalizes with DE-Cad in other tissues that require AJ-mediated adhesion. Dorsal closure (DC) is an embryonic process in which the migrating ectoderm closes the dorsal hole [54–56]. The amnioserosa, an extra-embryonic tissue, covers the dorsal hole and contributes to ectodermal closure by providing contractile forces that pull the contralateral ectodermal sheets together [54, 57, 58]. DC requires DE-Cad-mediated adhesion for ectodermal migration and fusion [59, 60]. To analyze Raskol and DE-Cad dynamics, we conducted time-lapse live imaging of embryos expressing YFP-tagged Raskol and RFP-tagged DE-Cad during DC. Colocalization of Raskol and DE-Cad was observed both at the amnioserosa cell contacts as well as in the dorsal most ectodermal cells at the zippering interface (S3A-D Fig). Raskol and DE-Cad colocalize in multiple *Drosophila* tissues, suggesting that Raskol may be a fundamental regulator of DE-Cad.

### Raskol regulates the distribution of polarized actin protrusions

Analysis of Raskol-YFP protein localization in BC clusters revealed that cytoplasmic levels of Raskol were ∼2x higher in PCs compared to BCs (S2B Fig and S4A Fig). Similarly, DE-Cad levels were up-regulated in PCs relative to BCs (Fig. 4A) [24] suggesting that Raskol protein expression pattern trends with DE-Cad. Accordingly, *shg* and raskol gene expression patterns overlap during embryonic development and both peak at 6–8hrs after egg laying (S4B Fig) [48]. To determine if Raskol is required for maintaining cell adhesions in PCs similar to BCs, we expressed *raskol* RNAi using *unpaired*-GAL4 (*upd*-GAL4) to drive expression specifically in the PCs [61]. Egg chambers were stained for F-actin (phalloidin) and nuclei (DAPI). Expression of control RNAi did not affect cluster adherence; however, *shg* RNAi expression caused cluster disassociation in ∼80% egg chambers (Fig 6A, 6B and 6D), similar to a previous report [24]. Expression of *raskol* RNAi in PCs caused BC disassociation in 63% of egg chambers (Fig 6C and 6D), suggesting that Raskol and DE-Cad might function together to promote BC cluster adhesion in both PCs and BCs.

**Fig 6.**
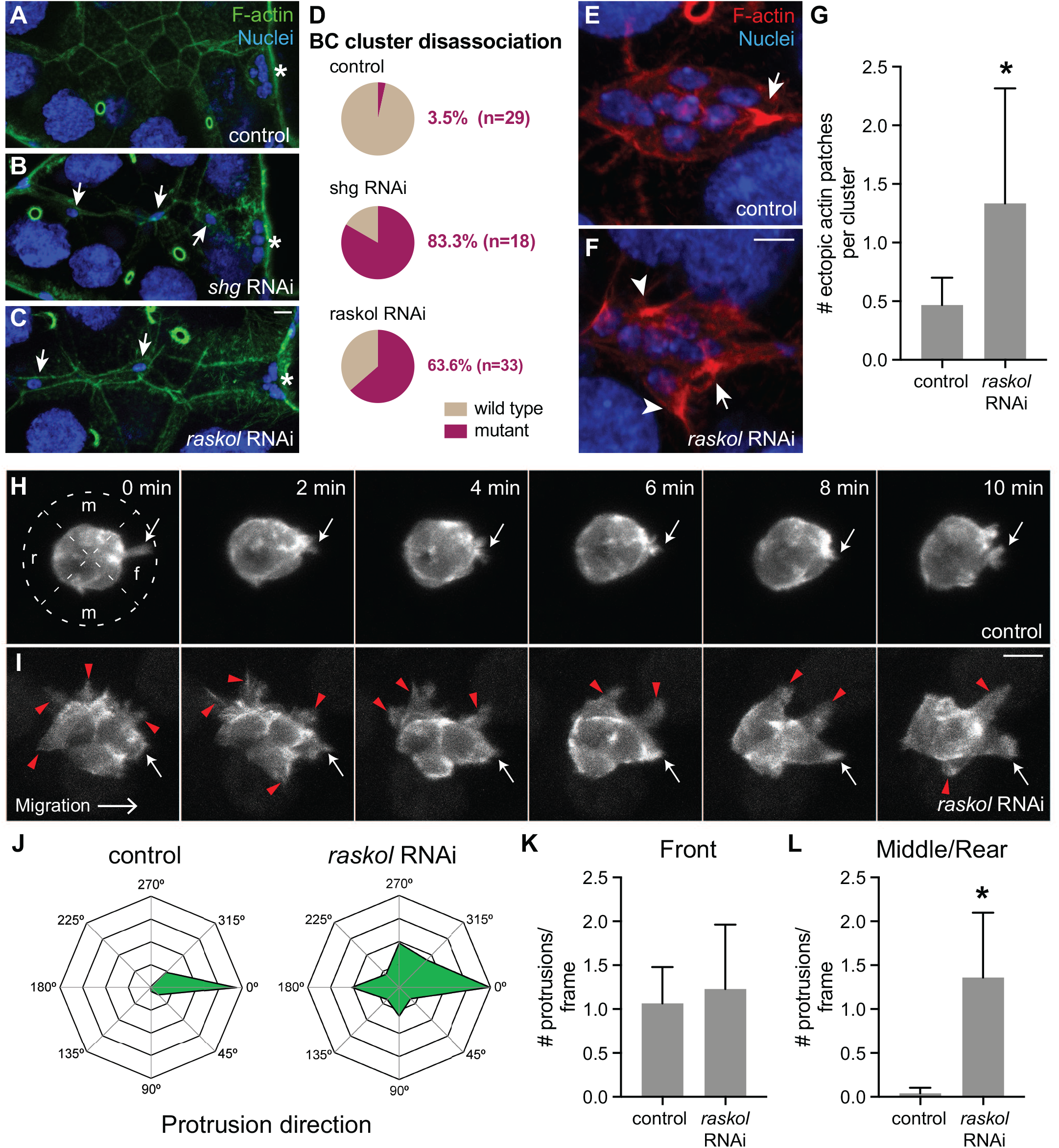
Raskol regulates actin organization in migrating BC clusters. A-C. Representative images of egg chambers stained for F-actin (green) and nuclei (blue) expressing control RNAi (A), *shg* RNAi (B) and *raskol* RNAi (C) in PCs using *upd*-GAL4. Asterisks mark the final position of the BC cluster adjacent to the oocyte. Arrows mark disassociated BC cells along the migratory path. Maximum projections of 10 μm stacks z-stacks are shown. D. Quantification of the cluster disassociation phenotype in control, *shg* and raskol RNAi-expressing BCs (p<0.0001). E-F. BC cluster expressing control RNAi (E) and raskol RNAi (F) under the control of slbo-GAL4 and stained for F-actin (red) and nuclei (blue). A white arrow marks the actin patch at the leading protrusion. White arrowheads mark ectopic actin patches around the BC cluster. G. Quantification of ectopic actin patches in control (n=30) and *raskol* (n=39) RNAi BC clusters (p<0.0001). H-I. Time-lapse images of migrating cluster expressing lifeact-GFP and control RNAi (H, n=8) or *raskol* RNAi (I, n=8) in BCs. White arrows mark the leading protrusion. Red triangles mark ectopic protrusions. f – front, m – middle and r – rear. J. Radar maps showing the distribution of protrusions around the BC cluster. 0° is the direction of migration. K. Number of front-oriented protrusions per frame observed in control and *raskol* RNAi expressing clusters. p=0.71. L. Number of protrusions per frame at the middle and rear of control RNAi and *raskol* RNAi-expressing clusters (p=0.0002). Scale bar in C is 10 μm and applies to A-C. Scale bar in panel F is 10 μm and applies to E-F. Scale bar in I is 10 μm and applies to H-I. Error bars in all graphs represent SD.

Next, we focused on the Raskol-YFP signal at the leading edge of the BC cluster (Fig 5B and 5D). Here, Raskol colocalized with actin suggesting that it may play a role in actin dynamics. Since Raskol contains a GAP domain and colocalizes with the cortical actin cytoskeleton in BCs, the FE, the amnioserosa and the dorsal most ectodermal cells [55, 60], we sought to determine whether it functions to regulate actin organization in BCs. We stained egg chambers expressing *raskol* RNAi in the BCs under the control of *slbo*-GAL4 driver for F-actin (phalloidin) and nuclei (DAPI) to assess F-actin distribution in the migrating cluster. In control BCs, actin accumulated at the base of protrusions typically oriented in the direction of migration (Fig 6E). In contrast, downregulation of raskol resulted in dramatic formation of multiple ectopic actin patches around the BC cluster (Fig 6F and 6G).

We then performed time-lapse live imaging of BC clusters expressing UAS-lifeact-GFP under the control of slbo-GAL4 to analyze protrusion dynamics in more detail. In control egg chambers, we observed protrusions extending primarily from the front of the migrating cluster (Fig 6H, 6J-L; Movie 1). In contrast, when *raskol* was downregulated, protrusions extended indiscriminately around the cluster (Fig 6I-L; Movie 2), consistent with our previous observations. Interestingly, the number of front-oriented protrusions in control and *raskol* RNAi expressing BCs did not differ significantly (Fig 6K). These data suggest that Raskol acts to restrict actin protrusions to the front of the BC cluster, which is critical to regulate BC migration. In addition, *raskol* knockdown caused BC delamination defects (Movie 3) and cluster disassociation (Movie 4) thereby confirming its importance in controlling cell adhesion and providing initial mechanistic insight into its role in regulating actin dynamics.

## Discussion

We combined evolution-guided bioinformatics with classical RNAi-based screening in *Drosophila* to identify regulators of DE-Cad-mediated cell adhesion. Our screen uncovered both established and novel regulators of DE-Cad function during BC migration. We demonstrated that one hit, the previously uncharacterized GAP domain containing protein Raskol, colocalizes with DE-Cad and regulates polarized actin dynamics in migrating BCs.

### ERC analysis reveals an evolutionary relationship between core components of the AJ

The core AJ components *shg, α-Cat* and *p120ctn* display high ERC values relative to one another, suggesting that the AJ complex has coevolved under selective pressure. α-Catenin provides the mechanical link between the cadherin-catenin complex and the actin cytoskeleton. Actin linkage is believed to be the original, ancestral function of the adherens junction that provided the foundation for multicellularity [9], so it is not surprising that *shg and α-Cat*, given their functional roles, have coevolved. Interestingly, p120-catenin is not essential for cadherin-mediated adhesion in flies [62], though it plays an important role in cadherin endocytosis in flies [63], similar to its established role in vertebrates [64]. Our ERC analysis suggests that shared selective pressures guided the evolution of the p120-catenin and DE-cadherin complex, and we speculate that these pressures may have shaped the range of p120-catenin functions in higher vertebrates.

Notably, no significant ERC relationship was observed between *shg* and *arm* or *shg* and other secondary AJ complex genes. While Arm is a core component of the AJ, it also functions as a key transcription factor in the Wnt signaling pathway [28, 29, 65]. We speculate that Arm function in Wnt signalling placed additional evolutionary pressures and altered its ERC signature relative to the other AJ genes. Similarly, neither *cno* nor vinc showed a strong ERC relationship to AJ genes, possibly reflecting their individual roles in AJ-independent processes [31, 66].

A number of genes identified in the *shg* ERC analysis have been implicated in regulating cell adhesion (Fig 1, Table 1). However, many of ERC-identified genes have not been functionally associated with DE-Cad, the AJ or cell adhesion. A previous genomic RNAi screen conducted in *Drosophila* identified multiple regulators of DE-Cad [43]. Notably, there was little overlap between the two screens. This highlights the potential of ERC analysis as an alternative, unbiased approach to generate a target gene list based solely on the evolutionary rate comparison. However, it is important to conduct secondary screens to validate the function of the target genes in a relevant biological system to eliminate false positives [16, 18, 19]. Also, ERC analysis cannot predict where (e.g., tissue type, cellular component) or when (e.g., developmental stage) a putative interaction will occur. Refinement of the *shg* ERC list based on spatio-temporal expression data can further eliminate false positive hits [19]. Nonetheless, a major advantage of ERC analysis is that genes that would otherwise not arise in a functional or associative genetic screen can be identified.

### DE-Cad and its regulators are required for BC migration

BC migration requires coordinated regulation of adhesion and motility [23, 67, 68] and is a good system for testing genes that regulate DE-Cad. BC migration is also an *in vivo* model for metastasis since many morphological characteristics of BCs resemble the invasive behaviour of metastatic cell clusters [68, 69]. However, in contrast to most models of epithelial-to-mesenchymal transition, the detachment of BC cluster requires upregulated levels of DE-Cad [23, 24]. DE-Cad-mediated adhesion between BCs and NCs is required for cluster polarization and directional migration, whereas adhesions between BCs and PCs are required for cluster adherence during migration [24, 25, 70]. Thus, BC migration is a useful system for genetic studies of cell adhesion and offers an opportunity to explore the role of adhesion genes in a relevant disease model [68, 69].

Our secondary genetic screen of *shg* ERC hits revealed potential roles for number of genes in BC migration, including kinases, GTPase regulators, transcription factors and cytoskeletal proteins. A small number of hits have putative or established roles in regulating DE-Cad and/or AJs. For example, PDZ-GEF was shown to colocalize with DE-Cad and function through GTPase Rap1 to regulate DE-Cad at the cell membrane [34, 35]. Hrb98DE is an RNA-binding protein that regulates DE-Cad mRNA processing [32, 33]. Mammalian orthologs of the small GTPase Rab5 regulate E-Cad trafficking [40, 42]. The mammalian homolog of transcription factor CG16952, Btbd7, regulates E-Cad expression [38, 39]. Additionally, genes such as RhoGAPpp190, *Stim, Appl* and *Rab5* were also identified in functional and proteomics screens of E-Cad [43, 44]. We also identified numerous genes that have not been linked to DE-Cad, including raskol, *CG11593, babo* and *zormin*. Out of these genes, *raskol*, CG11593 and *babo* directly regulate DE-Cad levels at the BC-BC contacts (Fig 4). Using *shg* ERC analysis and BC migration as a genetic model we have identified multiple novel genes that function to regulate DE-Cad-mediated cell adhesion in *Drosophila*.

### Raskol is a putative regulator of cell adhesion, polarity and actin dynamics

Downregulation of raskol caused severe BC cluster disassociation suggesting that Raskol is a critical regulator of BC adhesion. Consistent with this, Raskol colocalized with DE-Cad in multiple cell types and knockdown of *raskol* reduced DE-Cad levels at BC-BC cell contacts. Like *shg, raskol* is significantly upregulated in PCs relative to BCs and NCs. The encoded protein contains a highly conserved GAP domain that displays homology towards Ras- and Rho-GAPs, a plekstrin homology (PH) domain and a C2 domain that likely promote its membrane localization [71]. This suggests that, by colocalizing with DE-Cad, Raskol regulates adhesive strength between BCs to maintain cluster adhesion during detachment from the FE and subsequent migration. The mammalian orthologs of Raskol, Rasal2 and Dab2IP, were identified in a screen for RasGAP tumour suppressors [71] and are frequently downregulated in multiple types of cancer cells [53, 72–75]. Rasal2 and Dab2IP are capable of inactivating Ras through inducing GTP hydrolysis through their GAP domain and their downregulation leads to Ras overactivation [76–78]. Furthermore, inactivation of Rasal2 promotes invasive behaviour in a cell migration assay suggesting that Rasal2 has a conserved role in regulating cell adhesion and protrusive behaviour in mammals [71]. Dab2IP was identified in cadherin proximity biotinylation screens in mammalian epithelial cells [44] and mouse neonatal cardiomyocytes [79], further suggesting that the Rasal2/Dab2IP/Raskol family of proteins regulate AJ biology. Nonetheless, the mechanism of their function remains unclear.

Our study offers potential insight into Raskol function during collective migration. Epidermal growth factor receptor (EGFR) and PDGF- and VEGF-related receptor (PVR) localize to the leading edge of BC clusters and respond to a presumed gradient of guidance cues originating from the oocyte [67, 80–82]. The BC with the highest levels of EGFR/PVR activation becomes the leader cell and relays a signal to neighboring BCs through the DE-Cad adhesion complex to inhibit protrusion formation at the sides or rear of the cluster [24]. Interestingly, *gurken*, which encodes one of the four ligands for EGFR [81, 83, 84], also appeared on the *shg* ERC list (ERC value 0.62) as did its receptor *Egfr* (epidermal growth factor receptor, ERC value 0.36; S1 Table). The presence of both ligand and receptor suggests that EGF signaling has coevolved with DE-Cad to regulate cell adhesion. The primary GTPase that functions to regulate the directional migration of BC downstream of EGFR and PVR is Rac1, a member of the Rho GTPase family of proteins. We propose that Raskol, as a GAP, may function to suppress Rac1 signalling in non-leader BCs. Rac1 is expressed in all BCs, but the leader cells exhibit higher activity due to increased activation of EGFR and PVR [24, 80, 85]. Our results show that Raskol, like EGFR, PVR and Rac1 [80, 81, 85], restricts protrusions to the front of migrating BC cluster thus ensuring unidirectional migration. Downregulation of DE-Cad causes disruption in the polarized distribution of Rac1 in BC clusters, suggesting that DE-Cad regulates signaling downstream of EGFR and PVR [24, 86]. Therefore, Rac1 suppression might be achieved through Raskol GAP activity since knockdown of Rac1 or Raskol produce similar protrusion phenotypes [85]. Raskol may buffer the DE-Cad/Rac/actin mechanical feedback loop to regulate cell adhesion and promote collective cell migration. Whether Raskol directly interacts and regulates the GTPase domain of Rac1 remains to be explored.

Raskol localization is polarized with highest levels observed at the apical domain of ectodermal cells, FE cells and PCs. We also observe Raskol localization in the leading protrusion of BC cluster. This suggests that Raskol might regulate actin dynamics at the apical domain of polarized cells. However, Raskol does not directly regulate formation of protrusions since reducing Raskol levels does not affect the prevalence of leading protrusions. We predict that in leading BCs, Raskol limits active Rac1 to the tip of the protrusion to induce localized actin cytoskeletal remodelling. Accordingly, as reported by a Rac1-FRET sensor, high Rac1 activity is limited at the lamellopodial tip in the leading BC [24]. Overall, these data highlights two potential roles of Raskol function: 1) Raskol functions as a putative regulator of cell adhesion, and 2) Raskol regulates actin dynamics of the migrating cluster downstream of receptor tyrosine kinase signaling in BCs. Future studies dissecting the role of Raskol and other genes identified in our study are expected to offer insight in to the cooperative role of proteins that function along with the AJ to promote cell adhesion and cell migration.

## Methods and Materials

### Evolutionary Rate Covariance analysis

ERC values were calculated from protein coding sequences from 22 Drosophila species: *D. ananassae, D. biarmpies, D. bipectinada, D. elegans, D. erecta, D. eugracilis, D. ficusphila, D. grimshawi, D. kikawaii, D. persimilis, D. pseudoobscura, D. melanogaster, D. miranda, D. mojavensis, D. rhopaloa, D. sechelia, D. simulans, D. suzukii, D. takahashii, D. virilis, D. willistoni*, and *D. yakuba*. Protein coding sequences were downloaded from the Flybase website (http://www.flybase.org/) or the NCBI genome annotation website (https://www.ncbi.nlm.nih.gov/genome/annotation_euk/all/). Initially, coding sequences were evaluated for internal stop codons and the sequence was removed if found. For genes with multiple transcripts, the transcript with the longest sequence size was selected to represent the gene.

Orthology between genes across the multiple species were determined using the Orthofinder algorithm [87]. For each orthogroup, which are sets of genes that are orthologs and/or recent paralogs to each other, we omitted paralogous genes. Only orthogroups that had at least 6 species representation were analyzed further. Gene members of each orthogroup were aligned to each other using the PRANK aligner [88]. The multisequence alignment of each orthogroup was used by the PAML aaml program [89] to estimate the evolutionary rates on a single fixed species topology. A single species topology was estimated using a supertrees approach by combining individual orthogroup topologies that were estimated using RAxML [90]. Trees were combined using the matrix representation method implemented in phytools [91]

ERC was calculated using the branch lengths of each orthogroup. The overall species phylogenetic rates were normalized out for each orthogroup’s evolutionary rate, as described previously [18, 19]. Afterwards, ERC was measured as the Kendall’s τ correlation coefficients between two orthogroups and their species phylogeny normalized relative rates. ERC was then calculated for all pairwise orthogroup combinations.

### Gene ontology analysis

The *Drosophila* gene lists were subjected to GO analysis using Flybase website (http://www.flybase.org/). Homologs of *Drosophila* genes in mammalian genomes were generated using Flybase website (http://www.flybase.org/). A mammalian gene was considered a homolog if the gene was reported by 45% or more algorithms. Mammalian homologs were analyzed for canonical pathway and disease & function enrichment using Ingenuity Pathway Analysis tools (https://www.qiagenbioinformatics.com/products/ingenuity-pathway-analysis/).

### *Drosophila* melanogaster strains

All GAL4, reporter TRAP and RNAi stocks were obtained from Bloomington Drosophila Stock Center (BDSC) (S4 Table). *DE-Cad-GFP* and *DE-Cad-mCherry* knock-in stocks were used as E-Cad reporters [49]. s*lbo*-GAL4, UAS-lifeact-GFP, UAS-LacZ stock was generously provided by Jiong Chen (Nanjing University) [92]. Fly stocks were raised on standard yeast-based media at 20°C, unless otherwise noted.

### BC migration screen

For BC migration analysis, RNAi-expressing female flies under the control of *slbo-GAL4* were collected and transferred to vials containing fresh yeast paste and males. Flies were raised at 29ºC for 1–2 days. UAS-GFP was used as a reporter for RNAi expression. Dissected ovaries were fixed in 4% paraformaldehyde in PBS for 20 mins and washed 5 times with PBS. Ovaries were mounted on microscope slides in 70% glycerol and 20 μm z-stacks were acquired with a 20x objective on Nikon A1 scanning confocal microscope.

### Immunostaining of egg chambers

1–2 day old females were incubated at 29ºC for 1–2 days in vials with fresh yeast paste and males. Ovaries were dissected, fixed in 4% paraformaldehyde for 20 mins in PBS with 0.1% Triton-X (PBST), washed 5 times with PBST and blocked in normal goat serum (NGS) for 30 mins. For primary antibody staining, ovaries were incubated with Dcad2 antibody (1:100, Developmental Studies Hybridoma Bank) overnight at 4ºC and washed 10 times the next day over 1 hour. Next, ovaries were incubated with Alexa Fluor-conguated secondary antibody (1:150, Thermo Fisher Scientific) and Alexa 647 phalloidin (1:150, Thermo Fisher Scientific) for 2 hours. Egg chambers were then incubated in DAPI for 10 mins. Ovaries were washed 5 times in PBST and washed overnight and washed again 5 times next morning. Ovaries were stored and mounted on microscope slide in 70% glycerol and then imaged as described previously.

### Live imaging of BC clusters

Male flies containing slbo-GAL4, UAS-lifeact-GFP and UAS-LacZ were crossed to UAS-*raskol-RNAi* females. 1–2 day old F1 females were incubated at 29ºC for 1–2 days in vials with fresh yeast paste and *slbo-GAL4*, UAS-lifeact-GFP males. Ovaries were dissected for live imaging in imaging media (Schneiders’s medium, 15% fetal bovine serum (FBS) and 0.2mg/ml Insulin; Thermo Fisher Scientifc) according to published protocols [22, 93]. 100 μl of imaging media containing egg chambers was transferred to poly-D-lysine coated Mattek dishes for imaging. 20 μm Z-stacks (1 μm step size) covering the whole migrating border cell cluster were acquired every 2 minutes using a 40x objective on a Nikon A1 scanning confocal microscope.

### Live imaging of Raskol-YFP in embryos

Raskol-YFP [51] homozygous female flies were crossed to *DE-Cad-mCherry* [49] homozygous males. Embryos were collected overnight on grape juice agar plates and transferred to microscope slides coated with double-sided tape. Embryos were manually dechorionated and immediately transferred to halocarbon oil on coverslips with the dorsal side facing down. Coverslips were then attached to imaging chambers using double sided tape and imaged using a 60x objective on a Nikon A1 scanning confocal microscope.

### Quantification and statistics

Border cell migration defects were quantified as described previously [24]. Stage 10 egg chambers were analysed for each genotype. Border cell position along the migratory path was assigned into one of the following categories: 0–25% (no migration), 25–75% (delayed migration) and 75–100% (completed migration).

To quantify defects in border cell cluster adhesion, we determined the percentage of egg chambers where individual border cells had detached from the cluster.

To quantify DE-Cad levels, linescans across the BC-BC contacts were used to calculate the maximum pixel intensity at the contact in ImageJ. Peak values were then normalized to the peak intensity values of cell-cell contacts between NCs for each egg chamber. One-way ANOVA followed by Mann-Whitney tests were performed to determine significance. At least 22 border cell clusters (3 cell contacts per cluster) were imaged for each genotype.

To measure cytoplasmic levels of Raskol, an ROI was drawn in the cytoplasm of polar and border cells and average intensity determined. The ROI intensity was normalized to the average cytoplasmic intensity of Raskol-YFP in the nurse cells. To quantify colocalization of DE-Cad and Raskol, Pearson’s correlation coefficient (R) was calculated using the JACop plugin in ImageJ.

Ectopic actin patch number was quantified as described [92]. Actin patch present at the base of the leading edge protrusion was excluded from quantification since it is not ectopic. Welch’s t-test was performed to statistically compare number of ectopic actin patches between samples.

To quantify protrusion direction, we measured protrusions around the BC cluster in 45° increments at each frame of the movie (16 frames from 8 movies for each genotype). Protrusions between 315º and 45º angles were considered frontal protrusions; between 45º and 135º and 225º and 315º as middle protrusions; and between 225º and 135º as rear protrusions. Mann-Whitney test was performed to statistically compare number of protrusions between samples.

## Acknowledgements

We would like to thank Dr. Jiong Chen for providing slbo-lifeact-GFP stock and Bloomington Drosophila Stock Center for providing GAL4 and RNAi stocks. We also thank members of the Kwiatkowski lab for critical feedback on the manuscript.

## Movies

**Movie 1.** Border cell migration in control RNAi egg chambers. Lifeact-GFP and RNAi transgenes expressed under control of *slbo-GAL4. 30 mins*.

**Movie 2.** Border cell migration in *raskol* RNAi egg chambers.

**Movie 3.** Border cell delamination defects in *raskol* RNAi egg chambers.

**Movie 4.** Border cell cluster disassociation defects in *raskol RNAi egg chambers*.

## Supplemental Figure Legends

**S1 Fig.** DE-Cad levels are not reduced in BCs expressing RNAi against target genes with low ERC values. A-D. Representative images of BC clusters expressing DE-Cad-GFP and UAS-RNAi constructs in the BCs under the control of *slbo*-GAL4. Expression of UAS-RFP (A) and *CG5875* RNAi (D) did not result in significantly difference in DE-Cad levels at the BC-BC contacts relative to control. B. Expression of *Hrb98DE* RNAi (B) reduced DE-Cad levels at BC-BC contacts relative to control. C. Expression of capt RNAi (C) increased levels of DE-Cad at BC contacts relative to control. PC boundaries are marked by dashed lines. E. Quantification of DE-Cad levels at the BC-BC contacts of UAS-RFP (n=75), *Hrb98DE* (n=72), *capt* (n=60) and *CG5875* (n=60) RNAi expressing clusters. *** – p<0.001, ** – p<0.01. Error bars represent mean ± SD. Scale bar in D is 10 μm and applies to A-D.

**S2 Fig.** Raskol colocalizes with DE-Cad in the FE. A. Dorsal view of the FE in egg chamber expressing Raskol-YFP and stained for DE-Cad (red) and F-actin (blue). Raskol is enriched at the apical surface of the FE and colocalizes with DE-Cad and Factin. B. Cross-sectional images of the FE. Raskol colocalizes with DE-Cad and F-actin at the apical membrane of FE cells (arrows). Raskol also colocalizes with DE-Cad at PC contacts (arrowheads). In B, FE apical membrane faces the NCs. Individual channels correspond to the outlined box in the merged image. Scale bar in B is 10 μm and applies to A and B.

**S3 Fig.** Raskol colocalizes with DE-Cad in the amnioserosa and ectodermal cells during DC. A-D. Time-lapse images of Raskol-YFP and DE-Cad-RFP embryos during DC. Raskol colocalizes with DE-Cad at cell contacts in the amnioserosa (arrows). Raskol colocalizes with DE-Cad at the zippering interface of dorsal most ectodermal cells (arrowheads). Individual channels correspond to the outlined box in the merged image. Scale bar in D is 10 μm and applies to all panels.

**S4 Fig.** Coexpression pattern of *shg* and *raskol*. A. Mean cytoplasmic levels of Raskol in PCs and BCs relative to NCs. Cytoplasmic levels of Raskol are significantly higher in PCs compared to BCs according to Welch’s t-test (n=58, p<0.0001). B. *shg* and *raskol* expression patterns display similar trends during embryonic development. RNA-seq based expression data (obtained from www.flybase.org) in *Drosophila* embryos were plotted for *shg* and *raskol during embryonic stages (2 hr increments)*. Expression of both *shg* and *raskol* peaks at 6–8 hr after egg laying.

**S1 Table.** Top 500 shg ERC hits.

**S2 Table.** Enrichment analysis of shg ERC human orthologs: canonical pathways.

**S3 Table.** Enrichment analysis of shg ERC human orthologs: disease.

**S4 Table.** RNAi stocks used in this study.

**S5 Table.** Border cell migration and cluster disassociation data.

## References

1. Harris TJ, Tepass U. Adherens junctions: from molecules to morphogenesis. Nature reviews Molecular cell biology. 2010;11(7):502–14. doi: 10.1038/nrm2927. PubMed PMID: 20571587.

2. Mege RM, Ishiyama N. Integration of Cadherin Adhesion and Cytoskeleton at Adherens Junctions. Cold Spring Harb Perspect Biol. 2017; 9(5). Epub 2017/01/18. doi: 10.1101/cshperspect.a028738. PubMed PMID: 28096263.

3. Yap AS, Gomez GA, Parton RG. Adherens Junctions Revisualized: Organizing Cadherins as Nanoassemblies. Dev Cell. 2015;35(1):12–20. Epub 2015/10/16. doi: 10.1016/j.devcel.2015.09.012. PubMed PMID: 26460944.

4. Brunet T, King N. The Origin of Animal Multicellularity and Cell Differentiation. Dev Cell. 2017;43(2):124–40. Epub 2017/10/25. doi: 10.1016/j.devcel.2017.09.016. PubMed PMID: 29065305; PubMed Central PMCID: PMCPMC6089241.

5. Miller PW, Clarke DN, Weis WI, Lowe CJ, Nelson WJ. The evolutionary origin of epithelial cell-cell adhesion mechanisms. Curr Top Membr. 2013;72:267–311. Epub 2013/11/12. doi: 10.1016/B978-0-12-417027-8.00008-8. PubMed PMID: 24210433; PubMed Central PMCID: PMCPMC4118598.

6. Hulpiau P, Gul IS, van Roy F. New insights into the evolution of metazoan cadherins and catenins. Prog Mol Biol Transl Sci. 2013;116:71–94. Epub 2013/03/14. doi: 10.1016/B978-0-12-394311-8.00004-2. PubMed PMID: 23481191.

7. Miller PW, Pokutta S, Mitchell JM, Chodaparambil JV, Clarke DN, Nelson WJ, et al. Analysis of a vinculin homolog in a sponge (phylum Porifera) reveals that vertebrate-like cell adhesions emerged early in animal evolution. J Biol Chem. 2018;293(30):11674–86. Epub 2018/06/09. doi: 10.1074/jbc.RA117.001325. PubMed PMID: 29880641; PubMed Central PMCID: PMCPMC6066325.

8. Murray PS, Zaidel-Bar R. Pre-metazoan origins and evolution of the cadherin adhesome. Biol Open. 2014;3(12):1183–95. Epub 2014/11/15. doi: 10.1242/bio.20149761. PubMed PMID: 25395670; PubMed Central PMCID: PMCPMC4265756.

9. Rubsam M, Broussard JA, Wickstrom SA, Nekrasova O, Green KJ, Niessen CM. Adherens Junctions and Desmosomes Coordinate Mechanics and Signaling to Orchestrate Tissue Morphogenesis and Function: An Evolutionary Perspective. Cold Spring Harb Perspect Biol. 2017. Epub 2017/09/13. doi: 10.1101/cshperspect.a029207. PubMed PMID: 28893859.

10. Meng W, Takeichi M. Adherens junction: molecular architecture and regulation. Cold Spring Harb Perspect Biol. 2009;1(6):a002899. doi: 10.1101/cshperspect.a002899. PubMed PMID: 20457565; PubMed Central PMCID: PMC2882120.

11. Menke A, Giehl K. Regulation of adherens junctions by Rho GTPases and p120-catenin. Archives of biochemistry and biophysics. 2012;524(1):48–55. doi: 10.1016/j.abb.2012.04.019. PubMed PMID: 22583808.

12. Bertocchi C, Vaman Rao M, Zaidel-Bar R. Regulation of adherens junction dynamics by phosphorylation switches. Journal of signal transduction. 2012;2012:125295. doi: 10.1155/2012/125295. PubMed PMID: 22848810; PubMed Central PMCID: PMC3403498.

13. Padmanabhan A, Rao MV, Wu Y, Zaidel-Bar R. Jack of all trades: functional modularity in the adherens junction. Curr Opin Cell Biol. 2015;36:32–40. Epub 2015/07/21. doi: 10.1016/j.ceb.2015.06.008. PubMed PMID: 26189061.

14. Garcia MA, Nelson WJ, Chavez N. Cell-Cell Junctions Organize Structural and Signaling Networks. Cold Spring Harb Perspect Biol. 2018;10(4). Epub 2017/06/11. doi: 10.1101/cshperspect.a029181. PubMed PMID: 28600395; PubMed Central PMCID: PMCPMC5773398.

15. Clark NL, Alani E, Aquadro CF. Evolutionary rate covariation in meiotic proteins results from fluctuating evolutionary pressure in yeasts and mammals. Genetics. 2013;193(2):529–38. doi: 10.1534/genetics.112.145979. PubMed PMID: 23183665; PubMed Central PMCID: PMC3567741.

16. Findlay GD, Sitnik JL, Wang W, Aquadro CF, Clark NL, Wolfner MF. Evolutionary rate covariation identifies new members of a protein network required for Drosophila melanogaster female post-mating responses. PLoS genetics. 2014;10(1):e1004108. doi: 10.1371/journal.pgen.1004108. PubMed PMID: 24453993; PubMed Central PMCID: PMC3894160.

17. Wolfe NW, Clark NL. ERC analysis: web-based inference of gene function via evolutionary rate covariation. Bioinformatics. 2015;31(23):3835–7. doi: 10.1093/bioinformatics/btv454. PubMed PMID: 26243019; PubMed Central PMCID: PMC4751245.

18. Priedigkeit N, Wolfe N, Clark NL. Evolutionary signatures amongst disease genes permit novel methods for gene prioritization and construction of informative gene-based networks. PLoS genetics. 2015;11(2):e1004967. doi: 10.1371/journal.pgen.1004967. PubMed PMID: 25679399; PubMed Central PMCID: PMC4334549.

19. Clark NL, Alani E, Aquadro CF. Evolutionary rate covariation reveals shared functionality and coexpression of genes. Genome research. 2012;22(4):714–20. doi: 10.1101/gr.132647.111. PubMed PMID: 22287101; PubMed Central PMCID: PMC3317153.

20. Godin SK, Meslin C, Kabbinavar F, Bratton-Palmer DS, Hornack C, Mihalevic MJ, et al. Evolutionary and functional analysis of the invariant SWIM domain in the conserved Shu2/SWS1 protein family from Saccharomyces cerevisiae to Homo sapiens. Genetics. 2015;199(4):1023–33. doi: 10.1534/genetics.114.173518. PubMed PMID: 25659377; PubMed Central PMCID: PMC4391554.

21. Ziegler AB, Augustin H, Clark NL, Berthelot-Grosjean M, Simonnet MM, Steinert JR, et al. The Amino Acid Transporter JhI-21 Coevolves with Glutamate Receptors Impacts NMJ Physiology, and Influences Locomotor Activity in Drosophila Larvae. Scientific reports. 2016;6:19692. doi: 10.1038/srep19692. PubMed PMID: 26805723; PubMed Central PMCID: PMC4726445.

22. Prasad M, Wang X, He L, Cai D, Montell DJ. Border Cell Migration: A Model System for Live Imaging and Genetic Analysis of Collective Cell Movement. Methods in molecular biology. 2015;1328:89–97. doi: 10.1007/978-1-4939-2851-4_6. PubMed PMID: 26324431; PubMed Central PMCID: PMC4762686.

23. Montell DJ, Yoon WH, Starz-Gaiano M. Group choreography: mechanisms orchestrating the collective movement of border cells. Nature reviews Molecular cell biology. 2012;13(10):631–45. doi: 10.1038/nrm3433. PubMed PMID: 23000794; PubMed Central PMCID: PMC4099007.

24. Cai D, Chen SC, Prasad M, He L, Wang X, Choesmel-Cadamuro V, et al. Mechanical feedback through E-cadherin promotes direction sensing during collective cell migration. Cell. 2014;157(5):1146–59. doi: 10.1016/j.cell.2014.03.045. PubMed PMID: 24855950; PubMed Central PMCID: PMC4118667.

25. Niewiadomska P, Godt D, Tepass U. DE-Cadherin is required for intercellular motility during Drosophila oogenesis. The Journal of cell biology. 1999;144(3):533–47. PubMed PMID: 9971747; PubMed Central PMCID: PMC2132905.

26. Fulga TA, Rorth P. Invasive cell migration is initiated by guided growth of long cellular extensions. Nature cell biology. 2002;4(9):715–9. Epub 2002/08/29. doi: 10.1038/ncb848. PubMed PMID: 12198500.

27. Harris TJ. Adherens junction assembly and function in the Drosophila embryo. International review of cell and molecular biology. 2012;293:45–83. doi: 10.1016/B978-0-12-394304-0.00007-5. PubMed PMID: 22251558.

28. Mosimann C, Hausmann G, Basler K. Beta-catenin hits chromatin: regulation of Wnt target gene activation. Nature reviews Molecular cell biology. 2009;10(4):276–86. doi: 10.1038/nrm2654. PubMed PMID: 19305417.

29. Clevers H, Nusse R. Wnt/beta-catenin signaling and disease. Cell. 2012;149(6):1192–205. doi: 10.1016/j.cell.2012.05.012. PubMed PMID: 22682243.

30. Atherton P, Stutchbury B, Jethwa D, Ballestrem C. Mechanosensitive components of integrin adhesions: Role of vinculin. Experimental cell research. 2016;343(1):21–7. doi: 10.1016/j.yexcr.2015.11.017. PubMed PMID: 26607713; PubMed Central PMCID: PMC4856733.

31. Mandai K, Rikitake Y, Shimono Y, Takai Y. Afadin/AF-6 and canoe: roles in cell adhesion and beyond. Prog Mol Biol Transl Sci. 2013;116:433–54. doi: 10.1016/B978-0-12-394311-00019-4. PubMed PMID: 23481206.

32. Kourtidis A, Ngok SP, Pulimeno P, Feathers RW, Carpio LR, Baker TR, et al. Distinct E-cadherin-based complexes regulate cell behaviour through miRNA processing or Src and p120 catenin activity. Nature cell biology. 2015;17(9):1145–57. doi: 10.1038/ncb3227. PubMed PMID: 26302406; PubMed Central PMCID: PMC4975377.

33. Ji Y, Tulin AV. Poly (ADP-ribose) controls DE-cadherin-dependent stem cell maintenance and oocyte localization. Nature communications. 2012;3:760. doi: 10.1038/ncomms1759. PubMed PMID: 22453833; PubMed Central PMCID: PMC3319983.

34. Boettner B, Van Aelst L. The Rap GTPase activator Drosophila PDZ-GEF regulates cell shape in epithelial migration and morphogenesis. Molecular and cellular biology. 2007;27(22):7966–80. doi: 10.1128/MCB.01275-07. PubMed PMID: 17846121; PubMed Central PMCID: PMC2169160.

35. Spahn P, Ott A, Reuter R. The PDZ-GEF protein Dizzy regulates the establishment of adherens junctions required for ventral furrow formation in Drosophila. Journal of cell science. 2012;125(Pt 16):3801–12. doi: 10.1242/jcs.101196. PubMed PMID: 22553205.

36. Rudini N, Felici A, Giampietro C, Lampugnani M, Corada M, Swirsding K, et al. VE-cadherin is a critical endothelial regulator of TGF-beta signalling. The EMBO journal. 2008;27(7):993–1004. doi: 10.1038/emboj.2008.46. PubMed PMID: 18337748; PubMed Central PMCID: PMC2323269.

37. Fornetti J, Flanders KC, Henson PM, Tan AC, Borges VF, Schedin P. Mammary epithelial cell phagocytosis downstream of TGF-beta3 is characterized by adherens junction reorganization. Cell death and differentiation. 2016;23(2):185–96. doi: 10.1038/cdd.2015.82. PubMed PMID: 26113040; PubMed Central PMCID: PMC4716300.

38. Onodera T, Sakai T, Hsu JC, Matsumoto K, Chiorini JA, Yamada KM. Btbd7 regulates epithelial cell dynamics and branching morphogenesis. Science. 2010;329(5991):562–5. doi: 10.1126/science.1191880. PubMed PMID: 20671187; PubMed Central PMCID: PMC3412157.

39. Fan C, Miao Y, Zhang X, Liu D, Jiang G, Lin X, et al. Btbd7 contributes to reduced E-cadherin expression and predicts poor prognosis in non-small cell lung cancer. BMC cancer. 2014;14:704. doi: 10.1186/1471-2407-14-704. PubMed PMID: 25253020; PubMed Central PMCID: PMC4189533.

40. Yang J, Yao W, Qian G, Wei Z, Wu G, Wang G. Rab5-mediated VE-cadherin internalization regulates the barrier function of the lung microvascular endothelium. Cellular and molecular life sciences: CMLS. 2015;72(24):4849–66. doi: 10.1007/s00018-015-1973-4. PubMed PMID: 26112597; PubMed Central PMCID: PMC4827161.

41. Woichansky I, Beretta CA, Berns N, Riechmann V. Three mechanisms control Ecadherin localization to the zonula adherens. Nature communications. 2016;7:10834. doi: 10.1038/ncomms10834. PubMed PMID: 26960923; PubMed Central PMCID: PMC4792928.

42. Saitoh S, Maruyama T, Yako Y, Kajita M, Fujioka Y, Ohba Y, et al. Rab5-regulated endocytosis plays a crucial role in apical extrusion of transformed cells. Proceedings of the National Academy of Sciences of the United States of America. 2017;114(12):E2327–E36. doi: 10.1073/pnas.1602349114. PubMed PMID: 28270608; PubMed Central PMCID: PMC5373379.

43. Toret CP, D’Ambrosio MV, Vale RD, Simon MA, Nelson WJ. A genome-wide screen identifies conserved protein hubs required for cadherin-mediated cell-cell adhesion. The Journal of cell biology. 2014;204(2):265–79. doi: 10.1083/jcb.201306082. PubMed PMID: 24446484; PubMed Central PMCID: PMC3897182.

44. Guo Z, Neilson LJ, Zhong H, Murray PS, Zanivan S, Zaidel-Bar R. E-cadherin interactome complexity and robustness resolved by quantitative proteomics. Science signaling. 2014;7(354):rs7. doi: 10.1126/scisignal.2005473. PubMed PMID: 25468996; PubMed Central PMCID: PMC4972397.

45. Van Itallie CM, Tietgens AJ, Aponte A, Fredriksson K, Fanning AS, Gucek M, et al. Biotin ligase tagging identifies proteins proximal to E-cadherin, including lipoma preferred partner, a regulator of epithelial cell-cell and cell-substrate adhesion. Journal of cell science. 2014;127(Pt 4):885–95. Epub 2013/12/18. doi: 10.1242/jcs.140475. PubMed PMID: 24338363; PubMed Central PMCID: PMCPMC3924204.

46. Montell DJ, Rorth P, Spradling AC. slow border cells, a locus required for a developmentally regulated cell migration during oogenesis, encodes Drosophila C/EBP. Cell. 1992;71(1):51–62. PubMed PMID: 1394432.

47. Rorth P, Szabo K, Bailey A, Laverty T, Rehm J, Rubin GM, et al. Systematic gain-of-function genetics in Drosophila. Development. 1998;125(6):1049–57. PubMed PMID: 9463351.

48. Contrino S, Smith RN, Butano D, Carr A, Hu F, Lyne R, et al. modMine: flexible access to modENCODE data. Nucleic acids research. 2012;40(Database issue):D1082–8. doi: 10.1093/nar/gkr921. PubMed PMID: 22080565; PubMed Central PMCID: PMC3245176.

49. Huang J, Zhou W, Dong W, Watson AM, Hong Y. From the Cover: Directed, efficient, and versatile modifications of the Drosophila genome by genomic engineering. Proceedings of the National Academy of Sciences of the United States of America. 2009;106(20):8284–9. doi: 10.1073/pnas.0900641106. PubMed PMID: 19429710; PubMed Central PMCID: PMC2688891.

50. Cho KS, Lee JH, Kim S, Kim D, Koh H, Lee J, et al. Drosophila phosphoinositide-dependent kinase-1 regulates apoptosis and growth via the phosphoinositide 3-kinase-dependent signaling pathway. Proceedings of the National Academy of Sciences of the United States of America. 2001;98(11):6144–9. doi: 10.1073/pnas.101596998. PubMed PMID: 11344272; PubMed Central PMCID: PMC33436.

51. Lye CM, Naylor HW, Sanson B. Subcellular localisations of the CPTI collection of YFP-tagged proteins in Drosophila embryos. Development. 2014;141(20):4006–17. doi: 10.1242/dev.111310. PubMed PMID: 25294944; PubMed Central PMCID: PMC4197698.

52. Thul PJ, Akesson L, Wiking M, Mahdessian D, Geladaki A, Ait Blal H, et al. A subcellular map of the human proteome. Science. 2017;356(6340). doi: 10.1126/science.aal3321. PubMed PMID: 28495876.

53. Jia Z, Liu W, Gong L, Xiao Z. Downregulation of RASAL2 promotes the proliferation, epithelial-mesenchymal transition and metastasis of colorectal cancer cells. Oncology letters. 2017;13(3):1379–85. doi: 10.3892/ol.2017.5581. PubMed PMID: 28454265; PubMed Central PMCID: PMC5403410.

54. Hayes P, Solon J. Drosophila dorsal closure: An orchestra of forces to zip shut the embryo. Mechanisms of development. 2017;144(Pt A):2–10. doi: 10.1016/j.mod.2016.12.005. PubMed PMID: 28077304.

55. Kiehart DP, Crawford JM, Aristotelous A, Venakides S, Edwards GS. Cell Sheet Morphogenesis: Dorsal Closure in Drosophila melanogaster as a Model System. Annual review of cell and developmental biology. 2017;33:169–202. doi: 10.1146/annurevcellbio-111315-125357. PubMed PMID: 28992442.

56. Heisenberg CP. Dorsal closure in Drosophila: cells cannot get out of the tight spot. BioEssays: news and reviews in molecular, cellular and developmental biology. 2009;31(12):1284–7. doi: 10.1002/bies.200900109. PubMed PMID: 19882683.

57. Kiehart DP, Galbraith CG, Edwards KA, Rickoll WL, Montague RA. Multiple forces contribute to cell sheet morphogenesis for dorsal closure in Drosophila. The Journal of cell biology. 2000;149(2):471–90. PubMed PMID: 10769037; PubMed Central PMCID: PMC2175161.

58. Lynch HE, Crews SM, Rosenthal B, Kim E, Gish R, Echiverri K, et al. Cellular mechanics of germ band retraction in Drosophila. Developmental biology. 2013;384(2):205–13. doi: 10.1016/j.ydbio.2013.10.005. PubMed PMID: 24135149; PubMed Central PMCID: PMC3856716.

59. Gorfinkiel N, Arias AM. Requirements for adherens junction components in the interaction between epithelial tissues during dorsal closure in Drosophila. Journal of cell science. 2007;120(18):3289–98. doi: 10.1242/jcs.010850. PubMed PMID: WOS:000249559400015.

60. Duque J, Gorfinkiel N. Integration of actomyosin contractility with cell-cell adhesion during dorsal closure. Development. 2016;143(24):4676–86. doi: 10.1242/dev.136127. PubMed PMID: WOS:000393454600012.

61. Beccari S, Teixeira L, Rorth P. The JAK/STAT pathway is required for border cell migration during Drosophila oogenesis. Mechanisms of development. 2002;111(1–2):115–23. Epub 2002/01/24x. PubMed PMID: 11804783.

62. Myster SH, Cavallo R, Anderson CT, Fox DT, Peifer M. Drosophila p120catenin plays a supporting role in cell adhesion but is not an essential adherens junction component. Journal of Cell Biology. 2003;160(3):433–49. doi: 10.1083/jcb.200211083. PubMed PMID: WOS:000180840800017.

63. Bulgakova NA, Brown NH. Drosophila p120-catenin is crucial for endocytosis of the dynamic E-cadherin-Bazooka complex. Journal of cell science. 2016;129(3):477–82. doi: 10.1242/jcs.177527. PubMed PMID: WOS:000369505600004.

64. Kourtidis A, Ngok SP, Anastasiadis PZ. p120 catenin: an essential regulator of cadherin stability, adhesion-induced signaling, and cancer progression. Prog Mol Biol Transl Sci. 2013;116:409–32. Epub 2013/03/14. doi: 10.1016/B978-0-12-394311-8.00018-2. PubMed PMID: 23481205; PubMed Central PMCID: PMCPMC4960658.

65. Moon RT, Kohn AD, De Ferrari GV, Kaykas A. WNT and beta-catenin signalling: diseases and therapies. Nature reviews Genetics. 2004;5(9):691–701. doi: 10.1038/nrg1427. PubMed PMID: 15372092.

66. Bays JL, DeMali KA. Vinculin in cell-cell and cell-matrix adhesions. Cellular and molecular life sciences: CMLS. 2017;74(16):2999–3009. Epub 2017/04/13. doi: 10.1007/s00018-017-2511-3. PubMed PMID: 28401269; PubMed Central PMCID: PMCPMC5501900.

67. Cai D, Dai W, Prasad M, Luo J, Gov NS, Montell DJ. Modeling and analysis of collective cell migration in an in vivo three-dimensional environment. Proceedings of the National Academy of Sciences of the United States of America. 2016;113(15):E2134–41. doi: 10.1073/pnas.1522656113. PubMed PMID: 27035964; PubMed Central PMCID: PMC4839456.

68. Friedl P, Gilmour D. Collective cell migration in morphogenesis, regeneration and cancer. Nature reviews Molecular cell biology. 2009;10(7):445–57. doi: 10.1038/nrm2720. PubMed PMID: 19546857.

69. Stuelten CH, Parent CA, Montell DJ. Cell motility in cancer invasion and metastasis: insights from simple model organisms. Nature reviews Cancer. 2018;18(5):296–312. doi: 10.1038/nrc.2018.15. PubMed PMID: 29546880.

70. Geisbrecht ER, Sawant K, Su Y, Liu ZC, Silver DL, Burtscher A, et al. Genetic interaction screens identify a role for hedgehog signaling in Drosophila border cell migration. Developmental dynamics: an official publication of the American Association of Anatomists. 2013;242(5):414–31. doi: 10.1002/dvdy.23926. PubMed PMID: 23335293; PubMed Central PMCID: PMC3721345.

71. McLaughlin SK, Olsen SN, Dake B, De Raedt T, Lim E, Bronson RT, et al. The RasGAP gene, RASAL2, is a tumor and metastasis suppressor. Cancer cell. 2013;24(3):365–78. doi: 10.1016/j.ccr.2013.08.004. PubMed PMID: 24029233; PubMed Central PMCID: PMC3822334.

72. Sun L, Yao Y, Lu T, Shang Z, Zhan S, Shi W, et al. DAB2IP Downregulation Enhances the Proliferation and Metastasis of Human Gastric Cancer Cells by Derepressing the ERK1/2 Pathway. Gastroenterology research and practice. 2018;2018:2968252. doi: 10.1155/2018/2968252. PubMed PMID: 29743885; PubMed Central PMCID: PMC5884246.

73. Liu L, Xu C, Hsieh JT, Gong J, Xie D. DAB2IP in cancer. Oncotarget. 2016;7(4):3766–76. doi: 10.18632/oncotarget.6501. PubMed PMID: 26658103; PubMed Central PMCID: PMC4826168.

74. Dote H, Toyooka S, Tsukuda K, Yano M, Ouchida M, Doihara H, et al. Aberrant promoter methylation in human DAB2 interactive protein (hDAB2IP) gene in breast cancer. Clinical cancer research: an official journal of the American Association for Cancer Research. 2004;10(6):2082–9. PubMed PMID: 15041729.

75. Huang Y, Zhao M, Xu H, Wang K, Fu Z, Jiang Y, et al. RASAL2 down-regulation in ovarian cancer promotes epithelial-mesenchymal transition and metastasis. Oncotarget. 2014;5(16):6734–45. doi: 10.18632/oncotarget.2244. PubMed PMID: 25216515; PubMed Central PMCID: PMC4196159.

76. Hui K, Wu S, Yue Y, Gu Y, Guan B, Wang X, et al. RASAL2 inhibits tumor angiogenesis via p-AKT/ETS1 signaling in bladder cancer. Cellular signalling. 2018;48:38–44. doi: 10.1016/j.cellsig.2018.04.006. PubMed PMID: 29702203.

77. Noto S, Maeda T, Hattori S, Inazawa J, Imamura M, Asaka M, et al. A novel human RasGAP-like gene that maps within the prostate cancer susceptibility locus at chromosome 1q25. FEBS letters. 1998;441(1):127–31. PubMed PMID: 9877179.

78. Maertens O, Cichowski K. An expanding role for RAS GTPase activating proteins (RAS GAPs) in cancer. Advances in biological regulation. 2014;55:1–14. doi: 10.1016/j.jbior.2014.04.002. PubMed PMID: 24814062.

79. Li Y, Merkel CD, Yang X, Heier JA, Cantrell PS, Sun M, et al. The N-cadherin interactome in primary cardiomyocytes as defined by quantitative proximity proteomics. 2018. doi: 10.1101/348953%JbioRxiv.

80. Bianco A, Poukkula M, Cliffe A, Mathieu J, Luque CM, Fulga TA, et al. Two distinct modes of guidance signalling during collective migration of border cells. Nature. 2007;448(7151):362–5. doi: 10.1038/nature05965. PubMed PMID: 17637670.

81. McDonald JA, Pinheiro EM, Kadlec L, Schupbach T, Montell DJ. Multiple EGFR ligands participate in guiding migrating border cells. Developmental biology. 2006;296(1):94–103. doi: 10.1016/j.ydbio.2006.04.438. PubMed PMID: 16712835.

82. Duchek P, Somogyi K, Jekely G, Beccari S, Rorth P. Guidance of cell migration by the Drosophila PDGF/VEGF receptor. Cell. 2001;107(1):17–26. PubMed PMID: 11595182.

83. Goentoro LA, Reeves GT, Kowal CP, Martinelli L, Schupbach T, Shvartsman SY. Quantifying the Gurken morphogen gradient in Drosophila oogenesis. Dev Cell. 2006;11(2):263–72. doi: 10.1016/j.devcel.2006.07.004. PubMed PMID: 16890165; PubMed Central PMCID: PMC4091837.

84. Shilo BZ. Signaling by the Drosophila epidermal growth factor receptor pathway during development. Experimental cell research. 2003;284(1):140–9. PubMed PMID: 12648473.

85. Wang X, He L, Wu YI, Hahn KM, Montell DJ. Light-mediated activation reveals a key role for Rac in collective guidance of cell movement in vivo. Nature cell biology. 2010;12(6):591–7. doi: 10.1038/ncb2061. PubMed PMID: 20473296; PubMed Central PMCID: PMC2929827.

86. Ridley AJ. Rho GTPase signalling in cell migration. Curr Opin Cell Biol. 2015;36:103–12. doi: 10.1016/j.ceb.2015.08.005. PubMed PMID: 26363959; PubMed Central PMCID: PMC4728192.

87. Emms DM, Kelly S. OrthoFinder: solving fundamental biases in whole genome comparisons dramatically improves orthogroup inference accuracy. Genome biology. 2015;16:157. doi: 10.1186/s13059-015-0721-2. PubMed PMID: 26243257; PubMed Central PMCID: PMC4531804.

88. Loytynoja A, Goldman N. Phylogeny-aware gap placement prevents errors in sequence alignment and evolutionary analysis. Science. 2008;320(5883):1632–5. doi: 10.1126/science.1158395. PubMed PMID: 18566285.

89. Yang ZH. PAML 4: Phylogenetic analysis by maximum likelihood. Mol Biol Evol. 2007;24(8):1586–91. doi: 10.1093/molbev/msm088. PubMed PMID: WOS:000248848400003.

90. Stamatakis A. RAxML version 8: a tool for phylogenetic analysis and post-analysis of large phylogenies. Bioinformatics. 2014;30(9):1312–3. doi: 10.1093/bioinformatics/btu033. PubMed PMID: WOS:000336095100024.

91. Revell LJ. phytools: an R package for phylogenetic comparative biology (and other things). Methods Ecol Evol. 2012;3(2):217–23. doi: 10.1111/j.2041-210X.2011.00169.x. PubMed PMID: WOS:000302538500001.

92. Wang H, Qiu Z, Xu Z, Chen SJ, Luo J, Wang X, et al. aPKC is a key polarity determinant in coordinating the function of three distinct cell polarities during collective migration. Development. 2018;145(9). doi: 10.1242/dev.158444. PubMed PMID: 29636381.

93. Prasad M, Jang AC, Starz-Gaiano M, Melani M, Montell DJ. A protocol for culturing Drosophila melanogaster stage 9 egg chambers for live imaging. Nature protocols. 2007;2(10):2467–73. doi: 10.1038/nprot.2007.363. PubMed PMID: 17947988.

